# NEUROG3 Is Sufficient to Drive Neuroendocrine Differentiation in Prostate Cancer Cells

**DOI:** 10.1101/2025.11.10.687642

**Authors:** Abdelkader Daoud, Lu Han, Rachael K. Christensen, Jorge O. Múnera

**Affiliations:** Department of Regenerative Medicine and Cell Biology, Hollings Cancer Center, Medical University of South Carolina, Charleston, SC 29425; Department of Biochemistry and Molecular Biology, Hollings Cancer Center, Medical University of South Carolina, Charleston, SC 29425, USA

## Abstract

Treatment-emergent neuroendocrine prostate cancer (t-NEPC) arises following androgen deprivation therapy, leading to androgen-independent growth. Although multiple factors have been shown to be necessary for neuroendocrine differentiation (NED), their sufficiency has not been demonstrated. The prostate and colorectum share a common hindgut origin, and prostate neuroendocrine cell markers overlap with colorectal enteroendocrine cell (EEC) markers. Analysis of patient datasets revealed NEUROG3 amplification in both castration-resistant and neuroendocrine prostate cancers, correlating with poor survival. Because Neurogenin-3 (NEUROG3) is necessary and sufficient for EEC differentiation in the colorectum, we hypothesized that it could similarly drive NED in prostate cancer cells. A transient pulse of NEUROG3 repressed luminal identity and activated neuroendocrine programs, producing neuroendocrine cells within seven days. In summary, our findings identify NEUROG3 as a potential mediator of prostate cancer progression and establish a rapid *in vitro* model in which its transient activation is sufficient to initiate neuroendocrine differentiation.

## Introduction

Prostate cancer (PCa) is the second leading cause of cancer-related death in men in the United States. Androgen deprivation therapy (ADT) is the standard treatment for localized PCa, yet a subset of patients develops treatment-emergent neuroendocrine prostate cancer (t-NEPC)(1,2). Similarly, androgen withdrawal in the androgen-dependent LNCaP cell line induces neuroendocrine differentiation (NED), which reverses upon androgen restoration(3). Although multiple transcription factors have been shown to be required for NED(4-8), their sufficiency has typically depended on prolonged activation and additional stressors. Identifying a single factor capable of inducing NED under normal growth conditions would provide a valuable new model for studying NED.

The prostate and colorectum share a common developmental origin from the hindgut (9), and neuroendocrine cells (NECs) of the prostate closely resemble intestinal enteroendocrine cells (EECs) in both morphology and marker expression. Our comparative transcriptomic analyses show that the transcriptional signature of prostate NECs(10), resembles that of tracheal NECs, fetal pancreatic endocrine cells and large intestinal EECs. Neurogenin-3 (NEUROG3) is a pro-neurogenic transcription factor essential for endocrine cell specification throughout the gastrointestinal tract (11,12). NEUROG3 mRNA levels were significantly increased in castration-resistant NEPC compared to castration-resistant adenocarcinomas(13), and transcription factor analysis of t-NEPC versus prostate adenocarcinoma further identified NEUROG3 as a potential regulator of t-NEPC(14). In clinical data from prostate cancer patients, we found that amplification of NEUROG3 occurs in castration resistant prostate cancer (CRPC) and neuroendocrine prostate cancer (NEPC) compared to prostate adenocarcinoma, correlating with poor survival.

Transient NEUROG3 expression is sufficient to generate EECs in human pluripotent stem cell-derived gastric and colonic organoids(15,16). Based on the shared developmental origin of the prostate and colorectum and the similarities between NECs and EECs, we postulated that transient NEUROG3 expression could similarly induce NED in prostate cancer cells. To test this, we generated NEUROG3-deficient LNCaP cells through CRISPR-Cas9 mediated gene editing and we introduced a lentiviral doxycycline-inducible NEUROG3 construct to generate a NEUROG3-rescue cell line. Using this cell line, we demonstrate that transient activation of NEUROG3 is sufficient to inhibit the expression of AR and AR-responsive mRNAs enabling a subset of cells to undergo NED. The resulting NECs express synaptophysin and NEUROG3-induced cells acquire a transcriptional profile that resembles the neuroendocrine prostate cancer cell line NCI-H660. In summary, our study implicates NEUROG3 as an important factor in prostate cancer progression and provides a new *in vitro* system for studying NED.

## Materials and Methods

### Cell Culture

LNCaP (ATCC® CRL-1740™) and NCI-H660 (ATCC® CRL-5813™) prostate cancer cell lines were obtained from the American Type Culture Collection (ATCC, Manassas, VA, USA) and maintained under standard conditions. LNCaP cells were cultured in RPMI-1640 medium (Corning 10-041-CV) supplemented with 10 % fetal bovine serum (FBS; Gibco 26140-079) and 1× penicillin–streptomycin (100 U/mL penicillin and 100 µg/mL streptomycin; Gibco 15140-122). NCI-H660 cells were grown in RPMI-1640 medium (Corning 10-041-CV) supplemented with 5 % FBS, 5 µg/mL insulin (Sigma I9278), 5 µg/mL transferrin (Sigma T8158), 5 ng/mL sodium selenite (Sigma S5261), and 10 nM hydrocortisone (Sigma H0888). All cell lines were maintained at 37 °C in a humidified incubator with 5 % CO_2_ and were routinely tested for mycoplasma contamination.

### Generation of NEUROG3-Knockout Cells

NEUROG3-knockout (NEUROG3KO) LNCaP cells were generated using a CRISPR/Cas9 nuclease system following the same workflow previously established for generation of other knockout lines in the laboratory. A single-guide RNA targeting an early exon of NEUROG3 was cloned into the PX459 v2.0 vector (Addgene #62988). LNCaP cells were transfected with either PX459 alone or PX459-NEUROG3 using Lipofectamine 3000 (Thermo Fisher). Twenty-four hours after transfection, puromycin (5 µg/mL) was added for 48 hours to select transfected cells. Surviving cells were expanded and screened for NEUROG3 disruption by Sanger sequencing of the genomic region flanking the target site. Indel frequency and frameshift efficiency were quantified using Synthego ICE analysis.

### Generation of Doxycycline-Inducible NEUROG3 Rescue Cells

To generate doxycycline-inducible NEUROG3 rescue cells, full-length NEUROG3 cDNA was cloned into the pINDUCER20 lentiviral vector (Addgene #44012) through Gateway recombination using pDONR221 as an entry vector. Lentiviral particles were produced by co-transfecting 293T cells with pINDUCER20-NEUROG3 and the packaging plasmids psPAX2 (Addgene #12260) and pMD2.G (Addgene #12259) using Lipofectamine 3000. Viral supernatants were collected at 48 and 72 hours, filtered, and used to transduce NEUROG3KO LNCaP cells in the presence of 8 µg/mL polybrene. Infected cells were selected with G418 (500 µg/mL). To confirm inducibility, cells were treated with 1 µg/mL doxycycline for 24 hours and analyzed for NEUROG3 expression by immunofluorescence and western blotting.

### Immunofluorescence Staining

Cells were grown on 8 well IBIDI chamber slides and fixed with 4 % paraformaldehyde on ice for 15 min, permeabilized with 0.1 % Triton X-100, and blocked with 5 % bovine serum albumin. Primary antibodies against NEUROG3, synaptophysin (SYP), chromogranin A (CHGA), INSM1, NKX2-2, CDH1, and SCG2 were incubated overnight at 4 °C, followed by fluorescently conjugated secondary antibodies. Nuclei were counterstained with DAPI. Fluorescent images were acquired using a Zeiss LSM confocal microscope. The percentage of SYP-positive cells was quantified from at least three independent fields per condition using Imaris software.

### Western Blotting

Cells were lysed in Triton buffer containing 40 mM HEPES (pH 7.4), 150 mM NaCl, 2.5 mM MgCl_2_, and 1 % Triton X-100 supplemented with protease and phosphatase inhibitors. Lysates were cleared by centrifugation and quantified using the BCA assay. Equal protein amounts were separated by SDS-PAGE, transferred to PVDF membranes, and probed with antibodies against NEUROG3, CHGA, SYP, and β-actin. Bound antibodies were detected using HRP-conjugated secondary antibodies and enhanced chemiluminescence.

### RNA Sequencing and Analysis

Total RNA was extracted using the NucleoSpin RNA kit (Macherey-Nagel). RNA integrity was verified on an Agilent Bioanalyzer. Libraries were prepared using poly(A) selection and sequenced on an Illumina NovaSeq 6000 to yield approximately 300 million paired-end reads per sample. Reads were trimmed and aligned to hg38 using STAR, and expression values were quantified with RSEM. Differential expression analysis was performed with DESeq2. Gene Ontology enrichment was analyzed using ToppFun, and principal component analysis was conducted in Partek Flow. Heatmaps were generated in Python using log_2_(TPM + 1) values.

### Comparative Transcriptomic and Pathway Analyses

Transcriptomic profiles from doxycycline-induced NEUROG3 cells were compared with publicly available signatures for neuroendocrine prostate cancer and intestinal enteroendocrine lineages using ToppCell Atlas. Pathways enriched in vehicle-treated versus induced cells were identified using DESeq2 results, and ranked by adjusted p values. Principal component analysis confirmed that +DOX and vehicle-treated groups formed distinct clusters corresponding to luminal and neuroendocrine transcriptional states.

### Patient Data Analysis

Genomic and transcriptomic data for NEUROG3 in human prostate cancer were analyzed using normalized data from Laberecque *et al*. 2019(17) and using cBioPortal. Mutation, amplification, and deletion frequencies were retrieved from TCGA-PRAD and SU2C/PCF datasets. Kaplan–Meier survival analysis was performed to evaluate correlations between NEUROG3 expression and overall survival.

### Statistical Analysis

All experiments were performed in at least three experimental replicates. Data were analyzed using GraphPad Prism 9. Values are presented as mean ± SEM. Statistical significance was determined using unpaired two-tailed Student’s t-tests or one-way

ANOVA with Tukey’s post-hoc test where appropriate. P values < 0.05 were considered statistically significant.

## Results

### Prostate neuroendocrine cells and colon enteroendocrine cells share conserved features

NECs in normal prostate and prostate cancer are identified through immunostaining for the markers chromogranin A (CHGA) and synaptophysin (SYP). Immunofluorescence staining of normal human prostate revealed rare CHGA and SYP positive cells distributed within the E-CAD positive epithelium (Supplementary Figure S1A). Similar staining of normal human colon showed CHGA and SYP positive enteroendocrine cells located within glandular structures (Supplementary Figure S1A). Both NECs and EECs displayed a typical flask-like morphology. To determine whether these similarities extended beyond markers and morphology, we compared the transcriptional signature of prostate NECs defined by single-cell RNA-seq(10) with publicly available cell type atlases. ToppCell analysis revealed that the prostate neuroendocrine signature most closely matched NECs of the trachea, fetal pancreatic endocrine cells and large-intestinal EECs (Supplementary Figure S1B). These findings indicate that prostate NECs share morphology and key molecular features with EECs of the gastrointestinal tract.

### NEUROG3 amplification correlates with advanced disease and poor prognosis

*NEUROG3* mRNA levels are significantly increased in castration resistant neuroendocrine prostate cancer versus castration resistant adenocarcinomas(13). Furthermore, analysis of predicted transcription factors regulating the transcriptomic differences between t-NEPC and prostate adenocarcinoma identified NEUROG3 as a potential regulator (14). To further confirm the relevance of NEUROG3 to NEPC, we examined *NEUROG3* mRNA levels from a study that examined gene expression in metastatic castration-resistant prostate cancer (mCRPC) and identified 5 different molecular subtypes within the cohort (17). We found that *NEUROG3* levels were significantly elevated in a molecular subtype that represents NED (Figure 1A, S2A). These results indicate that increased *NEUROG3* expression correlates with NEPC across multiple studies.

**Figure 1.**
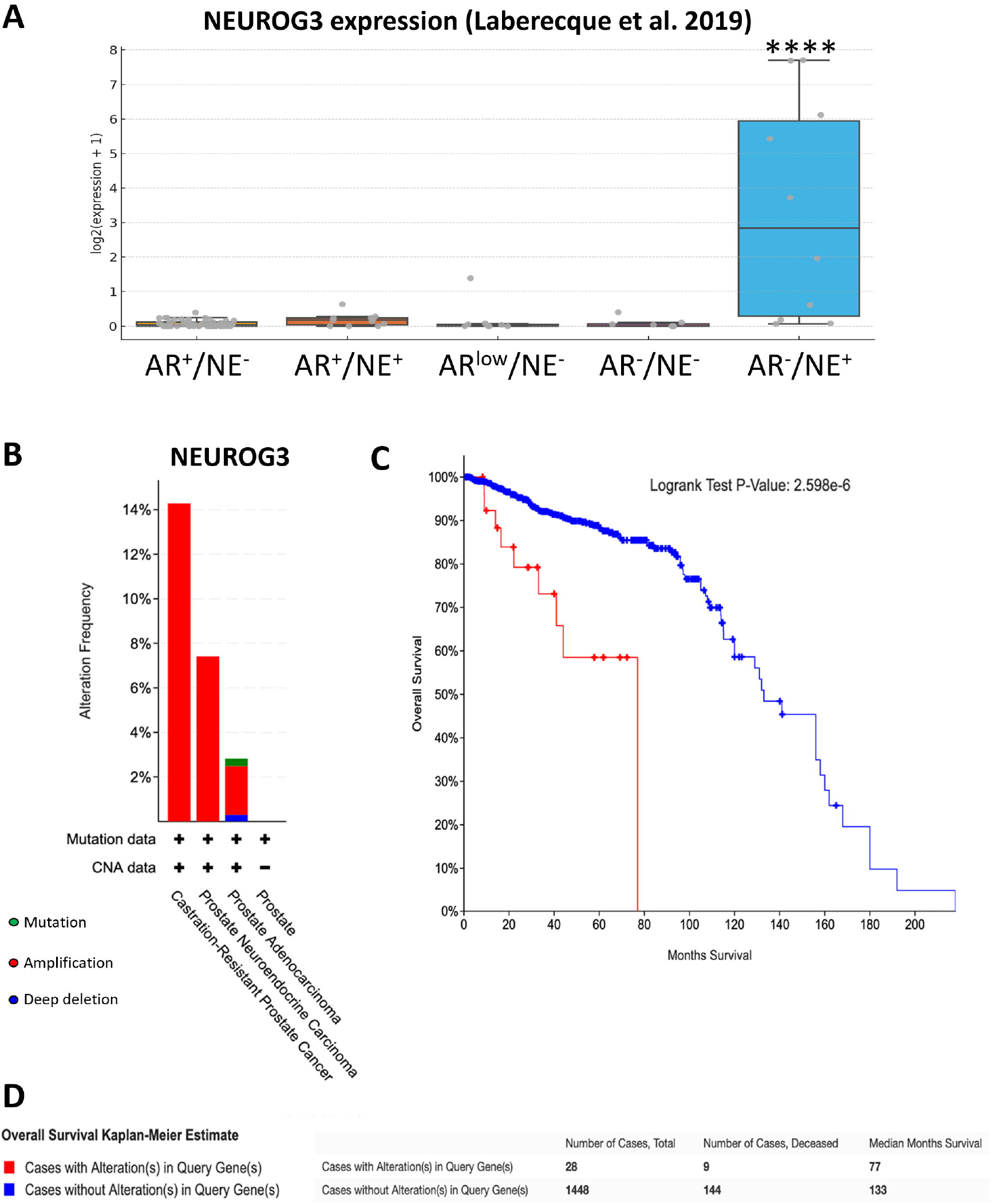
NEUROG3 alterations are associated with CRPC and decreased survival of patients. (A) NEUROG3 expression across molecular subtypes of metastatic castration-resistant prostate cancer (mCRPC), using transcript per million (TPM) data (log_2_[expression + 1]) from Labrecque et al. (2019). Subtypes include AR^+^/NE^−^, AR^+^/NE^+^, AR^low^/NE^−^, AR^−^/NE^−^, and AR^−^/NE^+^. NEUROG3 expression is significantly elevated in the AR^−^/NE^+^ subtype, with minimal expression observed in other subtypes. **** denotes p<0.0001. (B) Frequency of NEUROG3 gene alterations, including mutations, amplifications, and deep deletions, in prostate adenocarcinoma, prostate neuroendocrine carcinoma, and castration-resistant prostate cancer datasets from cBioPortal. (C) Kaplan–Meier analysis of overall survival in patients with or without NEUROG3 alterations. (D) Summary of overall survival data showing the number of cases, number of deceased patients, and median survival time for cohorts with and without NEUROG3 alterations.

To determine possible mechanisms by which *NEUROG3* mRNA expression is increased in NEPC, we analyzed patient data from cBioPortal. This analysis revealed that NEUROG3 is recurrently altered in prostate cancer, with amplification as the predominant event (Figure 1B). Alterations were most frequent in CRPCs and NEPCs compared with prostate adenocarcinomas. Kaplan–Meier analysis demonstrated that patients harboring NEUROG3 alterations had significantly reduced overall survival relative to unaltered cases (Figure 1C), and cohort summary data confirmed shorter median survival among patients with *NEUROG3* alterations (Figure 1D). These observations link *NEUROG3* amplification to aggressive disease behavior and poor survival.

### Development of an *in vitro* model of NEUROG3 mediated neuroendocrine differentiation

In mice, neurogenin-3 (*Ngn3*) is required for the generation of fetal pancreatic endocrine cells and EECs of the small and large intestine (11,12). Transient NEUROG3 expression is also sufficient to induce the differentiation of small and large intestinal EECs(15,16). Therefore, we predicted that transient NEUROG3 overexpression would be sufficient to induce NED in prostate cancer cells. Because NEUROG3 expression can be autoregulated(18), we generated *NEUROG3*KO cells using CRISPR-Cas9 mediated gene editing to minimize the effects of this autoregulation. Sequence analysis confirmed indels at the target site, and Synthego ICE deconvolution verified high editing efficiency and frameshift disruption resulting in premature stop codons (Supplementary Figure S3A). We introduced a lentiviral doxycycline-inducible *NEUROG3* transgene into the *NEUROG3*KO line thereby creating the LNCaP NEUROG3-rescue line (Supplementary Figure S3B). Immunofluorescence staining showed that NEUROG3 protein and its downstream effector NKX2-2 were robustly induced by doxycycline treatment but absent under vehicle conditions (Supplementary Figure S3C). Quantification of NEUROG3 positive nuclei confirmed efficient induction at day 1 and loss of expression by day 8 after doxycycline withdrawal (Supplementary Figure S3D). These data support that we generated a tightly regulated model to evaluate the temporal effects of transient NEUROG3 expression on LNCaP cell fate.

To determine if NEUROG3 could induce NED, we exposed LNCaP NEUROG3-rescue cells to vehicle (VEH) or doxycycline (+DOX) for 24 hours (day 1) followed by a change to regular growth media for 7 days (day 8). Western blotting revealed a significant induction of NEUROG3 protein at day 1 that subsided by day 8, whereas synaptophysin levels rose at day 8, indicating a delayed NED (Figure 2A). Densitometric quantification confirmed these changes in protein expression (Figure 2B). Immunofluorescence staining at day 8 showed that vehicle control cultures lacked SYP staining, whereas clusters of SYP-positive cells with neurite-like extensions emerged in the +DOX cultures (Figure 2C). Quantification demonstrated that approximately 8 percent of cells expressed synaptophysin after the pulse-chase compared with nearly zero percent in controls (Figure 2D). These results demonstrate that a short pulse of NEUROG3 expression was sufficient to initiate an NED program in a subset of prostate adenocarcinoma cells.

**Figure 2.**
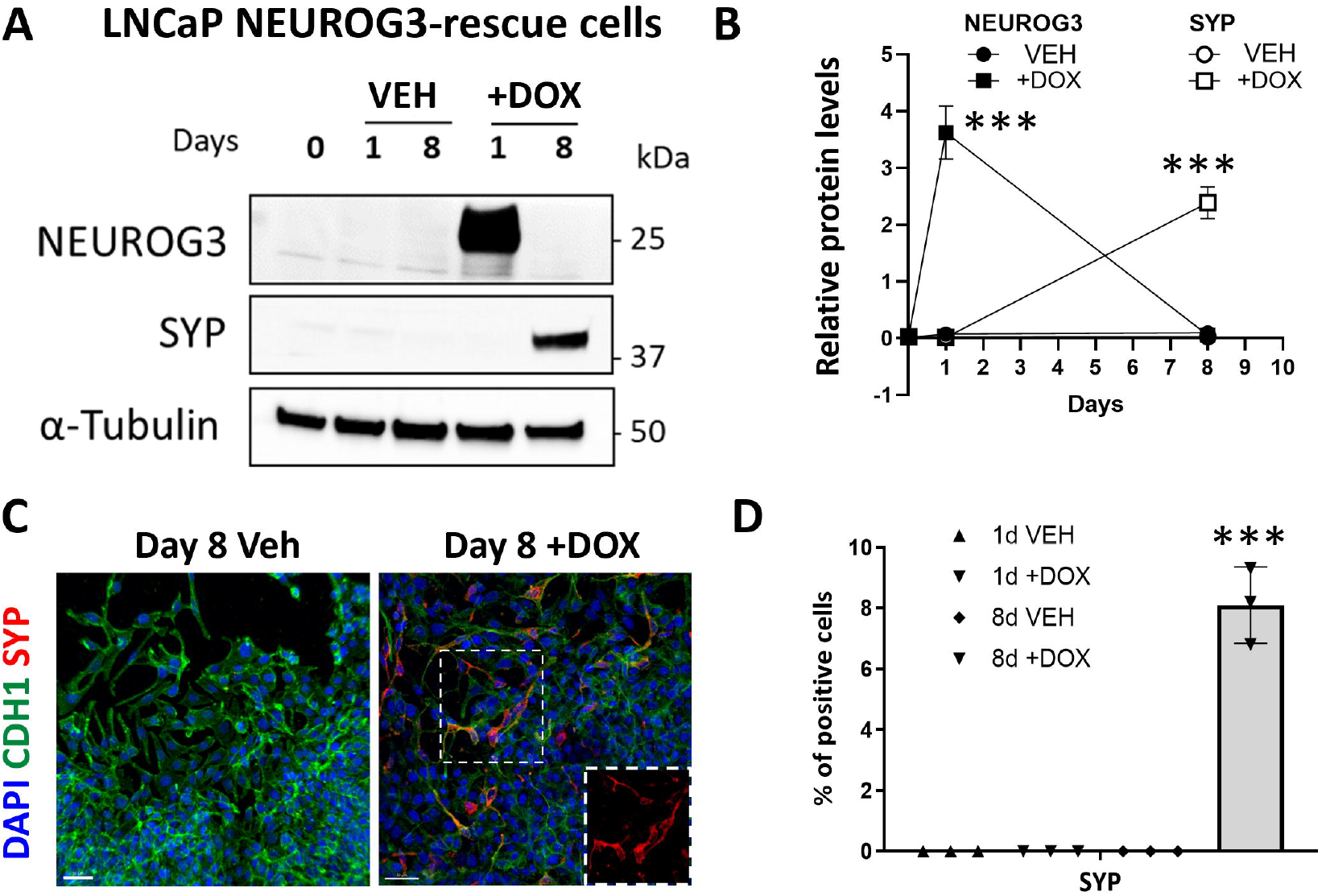
Transient NEUROG3 overexpression induces neuroendocrine differentiation of LNCaP cells. (A) Western blot analysis of NEUROG3, synaptophysin, and tubulin in LNCaP NEUROG3-rescue cells at day 0 (baseline), day 1 following a 24-hour pulse of vehicle or doxycycline, and day 8 (7-day chase following the 1-day pulse).(B) Densitometric quantification of NEUROG3 and synaptophysin normalized to tubulin across conditions. (C) Immunofluorescence staining for DAPI (blue), CDH1 (green), and synaptophysin (red) in day 8 vehicle- and doxycycline-treated cultures. Scale bars (D) Quantification of the percentage of synaptophysin-positive cells at day 1 and day 8 in vehicle- and doxycycline-treated cultures. Scale bars: (C) 50 µm.

### Transcriptomic validation of NEUROG3-induced differentiation

To define the transcriptional changes underlying NEUROG3-driven NED, we performed RNA-seq on LNCaP NEUROG3KO-rescue cells treated with vehicle (VEH) or doxycycline (+DOX) and compared them to the neuroendocrine prostate cancer line NCI-H660. Principal component analysis demonstrated that day 8 doxycycline-treated samples clustered more closely with NCI-H660 than day 8 vehicle controls, indicating partial transcriptional reprogramming toward a neuroendocrine phenotype (Figure 3A). Pathway enrichment analysis revealed that vehicle-treated samples were enriched for metabolic and hormone response pathways, including carboxylic acid and carbohydrate metabolism, while doxycycline-treated cells were enriched for neuronal programs such as synaptic signaling, neuron projection development, and synapse organization (Figure 3B). A heat map based on log_2_(TPM + 1) values showed coordinated upregulation of *NEUROG3* and its direct targets *NEUROD1, NEUROD2* and *NKX2-2* mRNAs after the 1-day pulse of doxycycline (Figure 3C). In addition to *AR*, luminal and AR-responsive mRNAs including *HOXB13, NKX3-1*, and *TMPRSS2*(19) were all downregulated following NEUROG3 induction. Transient NEUROG3 expression also induced the expression of mRNAs of multiple transcription factors that have been shown to be necessary for NED including *ASCL1*(6), *ONECUT2*(5,8) and *SOX2*(7). Interestingly, while *NEUROG3* expression returned to baseline levels at day 8, *ASCL1, ONECUT2* and *SOX2* mRNA levels remained elevated. Furthermore, neuroendocrine genes including *SYP, CHGA, INSM1*, and *SCG2* were highly induced on day 8. Overall, these results indicate that transient NEUROG3 induction activates a transcriptional cascade that durably upregulates key mediators of neuroendocrine differentiation ultimately driving the emergence of NECs on day 8.

**Figure 3.**
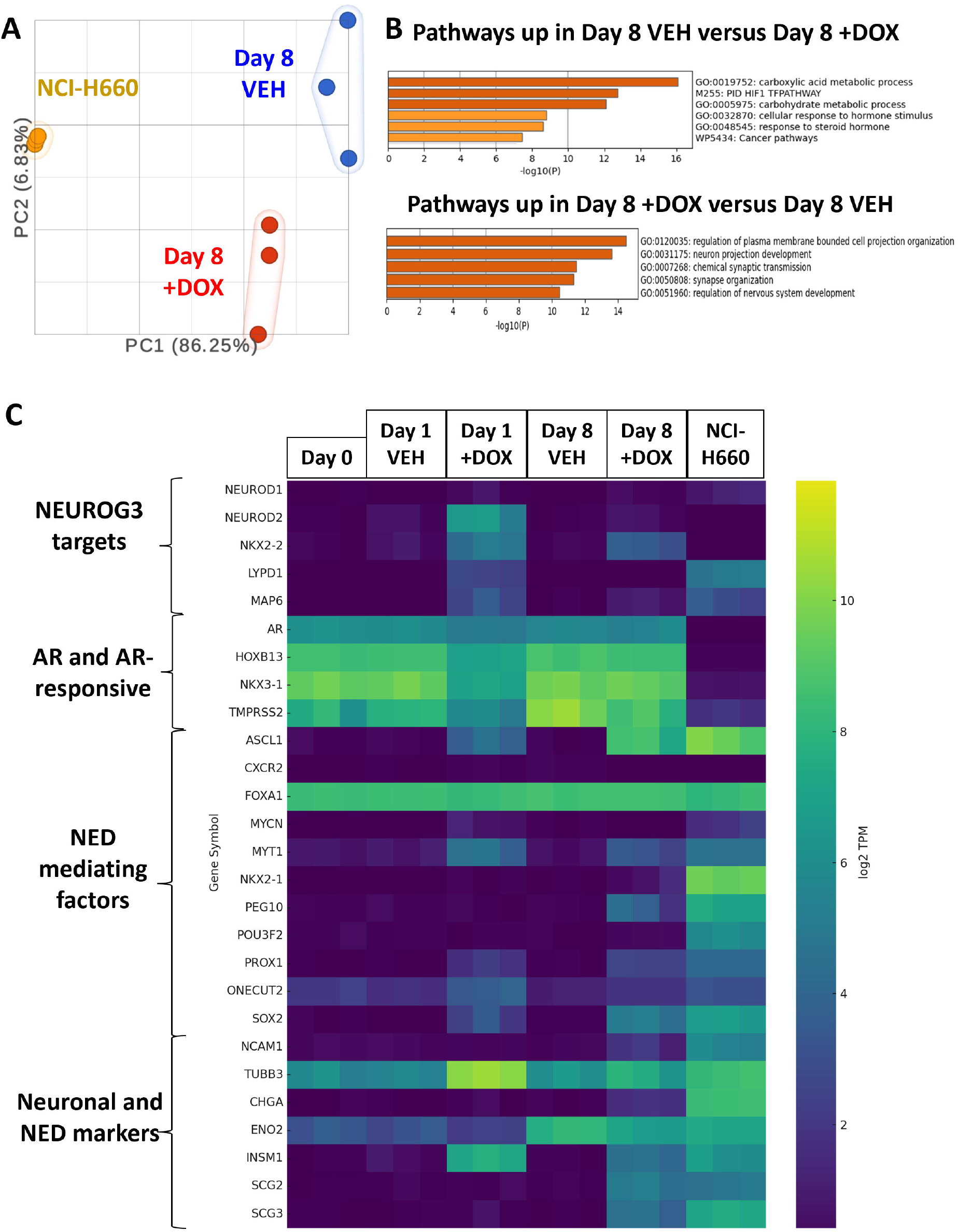
Transcriptomic analysis confirms neuroendocrine differentiation in NEUROG3 LNCaP cells. (A) Principal component analysis of RNA-seq data from day 8 vehicle-, day 8 doxycycline-treated LNCaP NEUROG3-rescue cells, and NCI-H660 cells. (B) Pathway enrichment analysis showing pathways upregulated in day 8 vehicle versus day 8 doxycycline and in day 8 doxycycline versus day 8 vehicle. (C) Heat map based on log2(TPM + 1) values showing expression of NEUROG3 and its targets, *AR* and AR-responsive genes, factors implicated in neuroendocrine differentiation, and neuronal/neuroendocrine cell markers.

## Discussion

Although multiple factors have been implicated in the process of neuroendocrine differentiation (NED), only a limited subset has demonstrated sufficiency, and even then, only under narrowly defined conditions. For example, constitutive expression of ONECUT2 has been shown to be sufficient to induce NED, but only under hypoxic conditions, which themselves are known to induce NEUROG3 and promote neuroendocrine lineage commitment(5,20). Likewise, ASCL1 has demonstrated the capacity to drive lineage conversion; however, this has primarily been observed in the 16DCRPC cell line, a model derived from LNCaP xenografts in castrated nude mice, where castration was performed post-tumor establishment(6). Notably, both ONECUT2 and ASCL1 require prolonged expression, further suggesting their roles may align more closely with the later phases of differentiation such as commitment or maturation rather than early cell fate specification.

In contrast, NEUROG3 appears to function as an upstream, transient developmental cue that initiates a cascade of neuroendocrine lineage programs in LNCaP cells. Unlike other transcription factors implicated in neuroendocrine differentiation (NED), which typically require sustained overexpression, NEUROG3 operates as a brief developmental pulse— 24 hours—yet induces a stable and progressive reprogramming of cell identity. Notably, in our model system, transient NEUROG3 induction leads to the activation of ASCL1 and ONECUT2, both of which remain persistently expressed through day eight, coinciding with the emergence of definitive neuroendocrine features. Thus, our results provide a mechanistic context for how ASCL1 and ONECUT2 fit within a broader developmental hierarchy, positioning NEUROG3 as an early trigger that initiates a neuroendocrine trajectory, which is subsequently reinforced and executed by sustained ASCL1 and ONECUT2 activity.

Although transient NEUROG3 induction produced neuroendocrine cells within days, only a subset of cells underwent complete reprogramming. This heterogeneity may reflect differences in NEUROG3 expression levels across the population, where only cells receiving an optimal dose of NEUROG3 achieve full lineage conversion. Alternatively, it may indicate that only a subset of LNCaP cells retains sufficient plasticity—perhaps a stem-like fraction capable of re-entering a developmental neuroendocrine trajectory. Future studies will address these possibilities by dissecting how NEUROG3 dosage and cellular state influence lineage outcomes. Moreover, while our data show sustained ASCL1 and ONECUT2 induction following transient NEUROG3 activation, additional work will be needed to determine whether these factors are directly regulated by NEUROG3 or act through downstream intermediates. Such studies will define the hierarchy of transcriptional events that couple transient NEUROG3 activation to durable neuroendocrine reprogramming.

## Acknowledgements

This work was funded by South Carolina Cancer Disparities Research Center (SC CADRE) NIH/NCI 5U54CA210962.

---

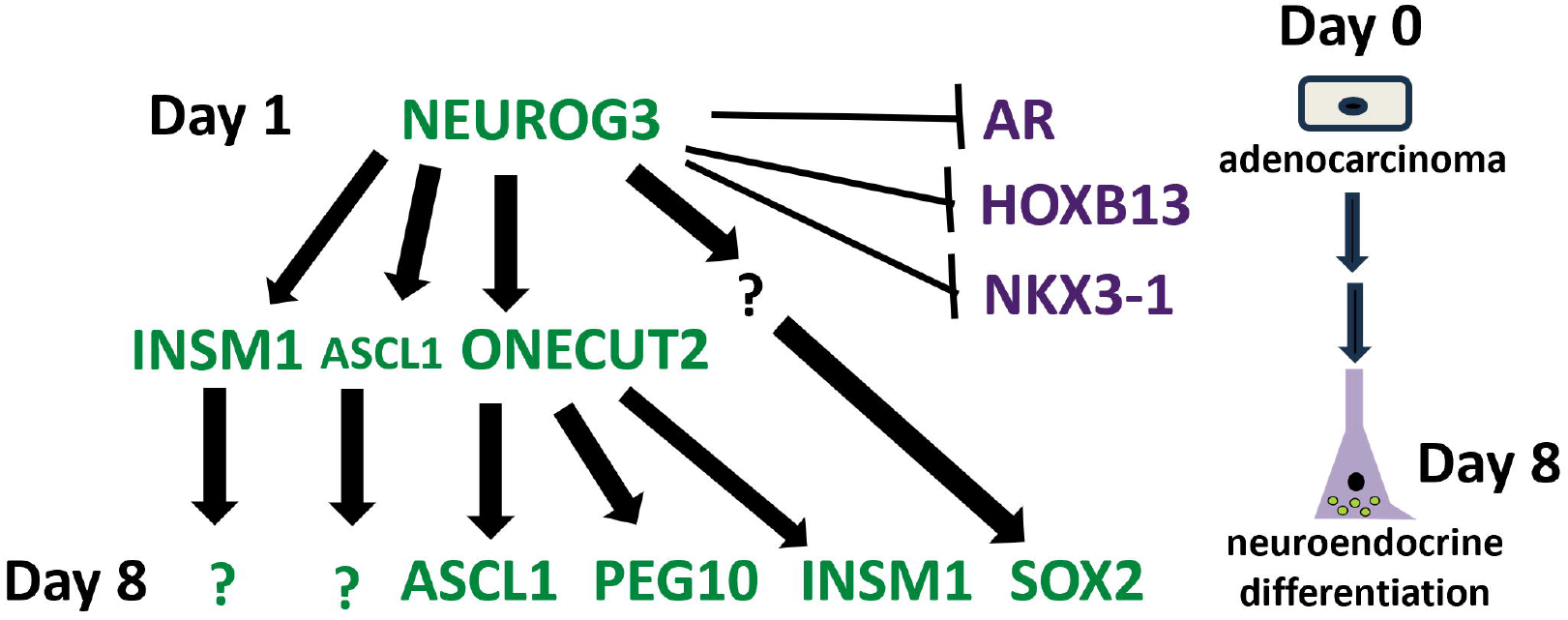

