## Supplementary information for "NEUROG3 Is Sufficient to Drive Neuroendocrine Differentiation in Prostate Cancer Cells"

**Figure S1. Prostate neuroendocrine cells resemble colon enteroendocrine cells.**

**A**

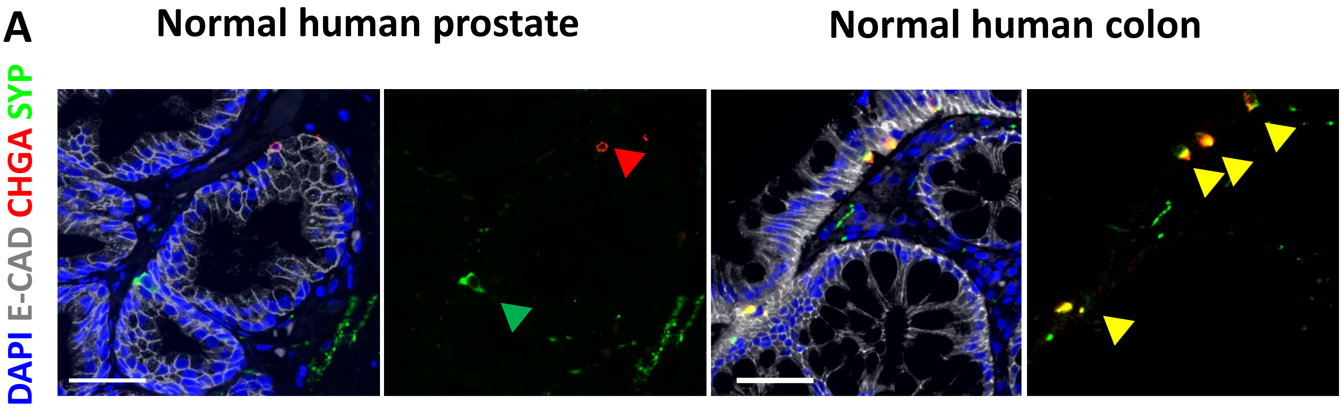


**B**

Prostate NEC signature from Henry *et al.* 2018

ToppCell Atlas

| Name | Source | pValue |
|---|---|---|
| Pulmonary NEC | Mouse Trachea Epithelial Cells (MTECs) | 3.17E-61 |
| **FetalPancreas-Fetal endocrine cell** | **Human Cell Landscape (Adult and Fetal)** | **1.47E-55** |
| Neuroendocrine\|World / Class top | Mouse Adult Trachea | 6.15E-49 |
| Large Intestine-Epithelial-glandular\| | Mouse Adult Overview - TabulaMuris | 1.08E-47 |
| **ts12 Large Intestine-Epithelial-glandular-Chromaffin Cell--enteroendocrine cell** | **Mouse Adult Overview - TabulaMuris** | **1.08E-47** |

### Figure S2. NEUROG3 Is Upregulated in a Neuroendocrine Subtype of metastatic Castration-Resistant Prostate Cancer

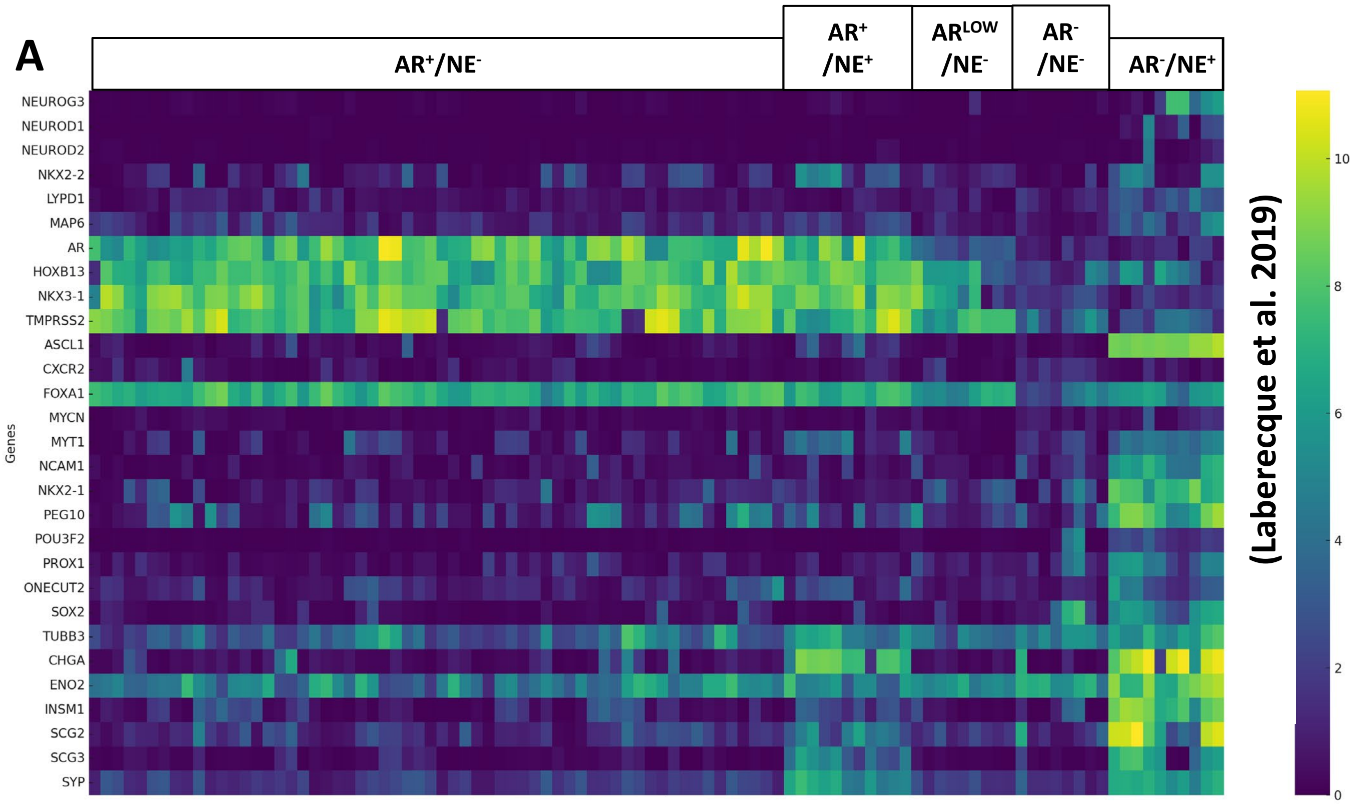

**Figure S3. Generation and validation of LNCaP NEUROG3KO and rescue cell line.**

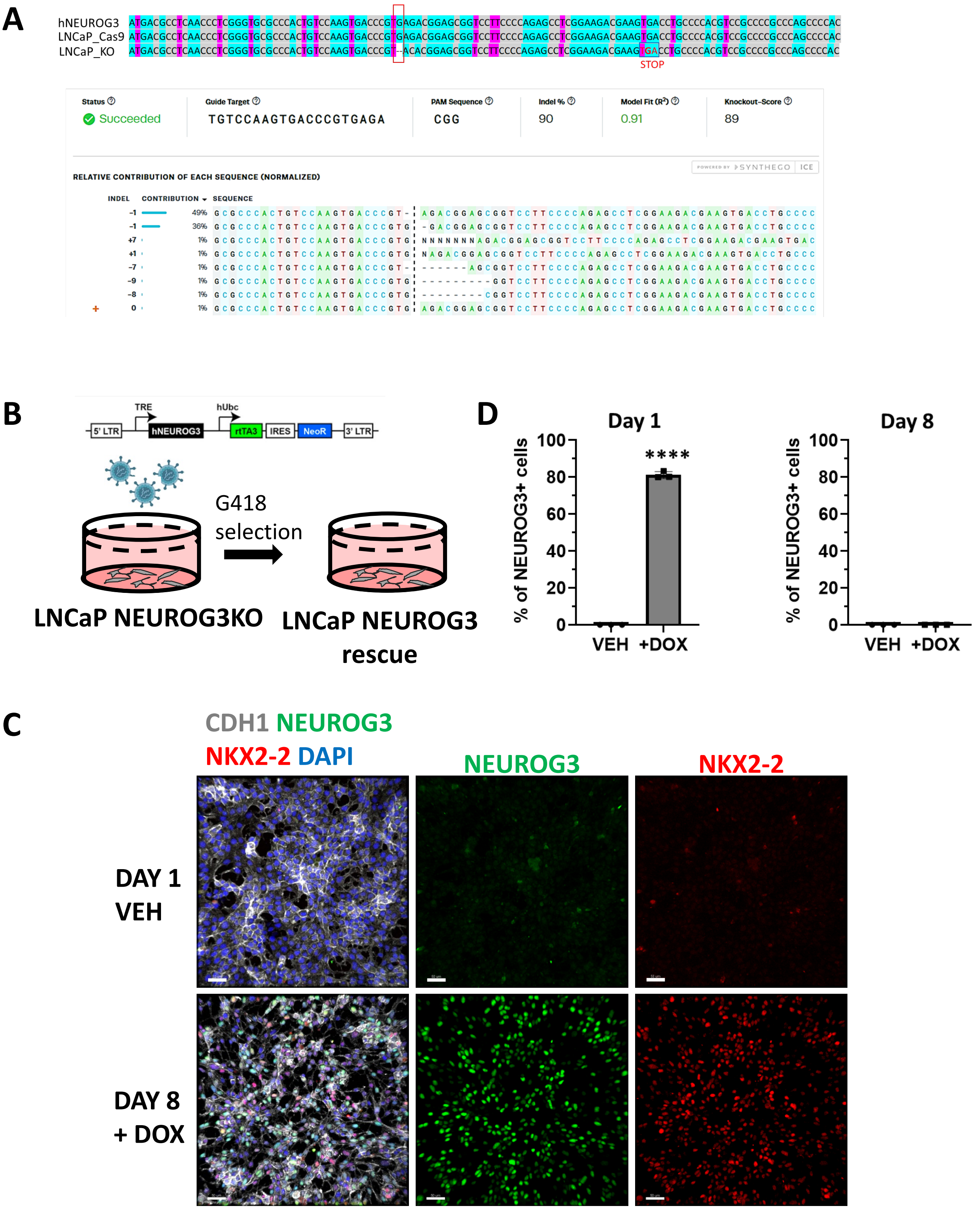

**Figure S1**: **Transcriptome of prostate neuroendocrine cells resembles fetal pancreatic endocrine cells and intestinal EECs.** (A) Immunofluorescence staining of normal human prostate tissue showing DAPI (blue), E-CAD (gray), CHGA (red), and SYP (green). Insets show CHGA- and SYP-positive cells indicated by arrowheads. (B) Immunofluorescence staining of normal human colon tissue showing DAPI (blue), E-CAD (gray), CHGA (red), and SYP (green). CHGA- and SYP-positive enteroendocrine cells are indicated by arrowheads. (C) ToppCell Atlas analysis showing enrichment of the prostate neuroendocrine carcinoma (NEC) gene signature from Henry et al. 2018 in fetal pancreatic endocrine cells and large intestine enteroendocrine cells. Scale bars: (A) 50 μm.

**Figure S2. NEUROG3 Is Upregulated in a Neuroendocrine Subtype of metastatic Castration-Resistant Prostate Cancer.** (A) Heat map based on log2(TPM + 1) values from RNA-seq data from Laberecque et al. 2019 showing expression of NEUROG3 and its targets, *AR* and AR-responsive genes, factors implicated in neuroendocrine differentiation, and neuronal/neuroendocrine cell markers.

**Figure S3**: **Generation and validation of NEUROG3-knockout and rescue LNCaP cells.** (A) Sequence alignment showing the CRISPR-Cas9–mediated NEUROG3-knockout in LNCaP cells with corresponding guide RNA target site, PAM sequence, and indel profile. Below, Synthego ICE analysis showing the distribution and relative contribution of indels in the NEUROG3 locus. (B) Schematic of doxycycline-inducible NEUROG3 rescue construct introduced into LNCaP NEUROG3 knockout cells. (C) Immunofluorescence staining for CDH1 (gray), NEUROG3 (green), NKX2-2 (red), and DAPI (blue) in vehicle- and doxycycline-treated LNCaP NEUROG3-rescue cells. (D) Quantification of NEUROG3-positive cells at day 1 and day 8 following treatment with vehicle or doxycycline. Scale bars: (C) 50 μm.
